# Chemically di-ubiquitylated H1 variants reveal position-dependent stimulation of the DNA repair protein RNF168

**DOI:** 10.1101/2024.07.19.604272

**Authors:** Pauline Franz, Charlotte M. Delvaux de Fenffe, Beat Fierz

## Abstract

Ubiquitylation of histone H2A at lysines 13 and 15 by the E3 ligase RNF168 plays a key role in orchestrating DNA double-strand break (DSB) repair, which is often deregulated in cancer. RNF168 activity is triggered by DSB signaling cascades, reportedly through K63-linked poly-ubiquitylation of linker histone H1. However, direct experimental evidence of this mechanism has been elusive, primarily due to the lack of methods to specifically poly-ubiquitylate H1. Here, we developed a versatile click-chemistry approach to covalently link multiple proteins in a site-specific, controlled, and stepwise manner. Applying this method, we synthesized H1 constructs bearing triazole-linked di-ubiquitin on four DNA repair-associated ubiquitylation hotspots (H1^Kx^Ub_2_, at K17, 46, 64 and 96). Integrated into nucleosome arrays, the H1^Kx^Ub_2_ variants stimulated H2A ubiquitylation by RNF168 in a position-dependent manner, with H1^K17^Ub_2_ showing the strongest RNF168 activation effect. Moreover, we show that di-ubiquitin binding is the driving force underlying RNF168 recruitment, introducing H1^K17^Ub_2_ into living U-2 OS cells. Together, our results support the hypothesis of poly-ubiquitylated H1 guiding RNF168 recruitment to DSB sites. Moreover, we demonstrate how the streamlined synthesis of H1^Kx^Ub_2_ variants enables mechanistic studies into RNF168 regulation, with potential implications for its inhibition in susceptible cancers.

## Introduction

DNA damage repair following DSBs involves signaling cascades of numerous regulatory proteins, which dynamically bind (‘read’) and alter (‘write’ or ‘erase’) post-translational modifications (PTMs) on histones, the main protein components of chromatin organizing the genome ^[1,2]^. The ubiquitin system, one of those PTM-based regulatory mechanisms, plays a key role in controlling the two major pathways of DSB repair, homologous recombination (HR) and non-homologous end-joining (NHEJ),^[3]^ both of which are deregulated in many diseases, especially in cancer ^[4,5]^. After detecting a DSB, poly-ubiquitin chains are installed on chromatin, marking the damage site (**Figure 1a**). This generates binding sites for the ubiquitin E3 ligase RNF168, a key protein that, when dysregulated in tumors, has been associated with genomic instability and disease progression.^[6]^ RNF168 acts as a ubiquitin ‘writer’ by installing H2AK13/15Ub.^[7]^ The placement of H2AK15Ub by RNF168 is essential for the DNA repair pathway choice ^[8]^ as it recruits two major downstream effectors, either 53BP1, promoting NHEJ,^[9,10]^ or the E3 ligase BRCA1/BARD1, initiating HR (**Figure 1a**).^[11–15]^ In addition, RNF168 acts as a ubiquitin ‘reader’ by binding to K63-linked poly-ubiquitin chains through its ubiquitin-dependent DSB recruitment module 1 (UDM1), as well as to H2AK13/15Ub through a second domain, UDM2 (**Figure 1a**).^[16–18]^ This ubiquitin-binding activity of RNF168, in particular to poly-ubiquitin chains via UDM1, is essential for its proper localization to DSB sites.^[19,20]^ Mutations that disrupt ubiquitin-binding by RNF168 result in disease, such as the RIDDLE syndrome.^[8,21–23]^ Intriguingly, the chromatin-associated protein that carries the initial poly-ubiquitin signal, serving as an anchoring point for RNF168, is still debated (**Figure 1a**). The linker histone H1, in particular subtype H1.2 (now generally referred to as H1), as well as another chromatin binding factor, L3MBTL3, have both been suggested as potential carriers.^[18]^ Moreover, mass spectrometry experiments identified several lysines in H1 that are ubiquitylated during DNA damage repair, strengthening the hypothesis of H1 playing a role (**Figure 1b**).^[24]^ However, whether the direct recognition of poly-ubiquitylated H1 by RNF168 controls its downstream activity remains a long-standing question in the DNA damage field (**Figure 1a**), also due to a lack of suitably modified H1 proteins that would allow mechanistic studies.

**Figure 1.**
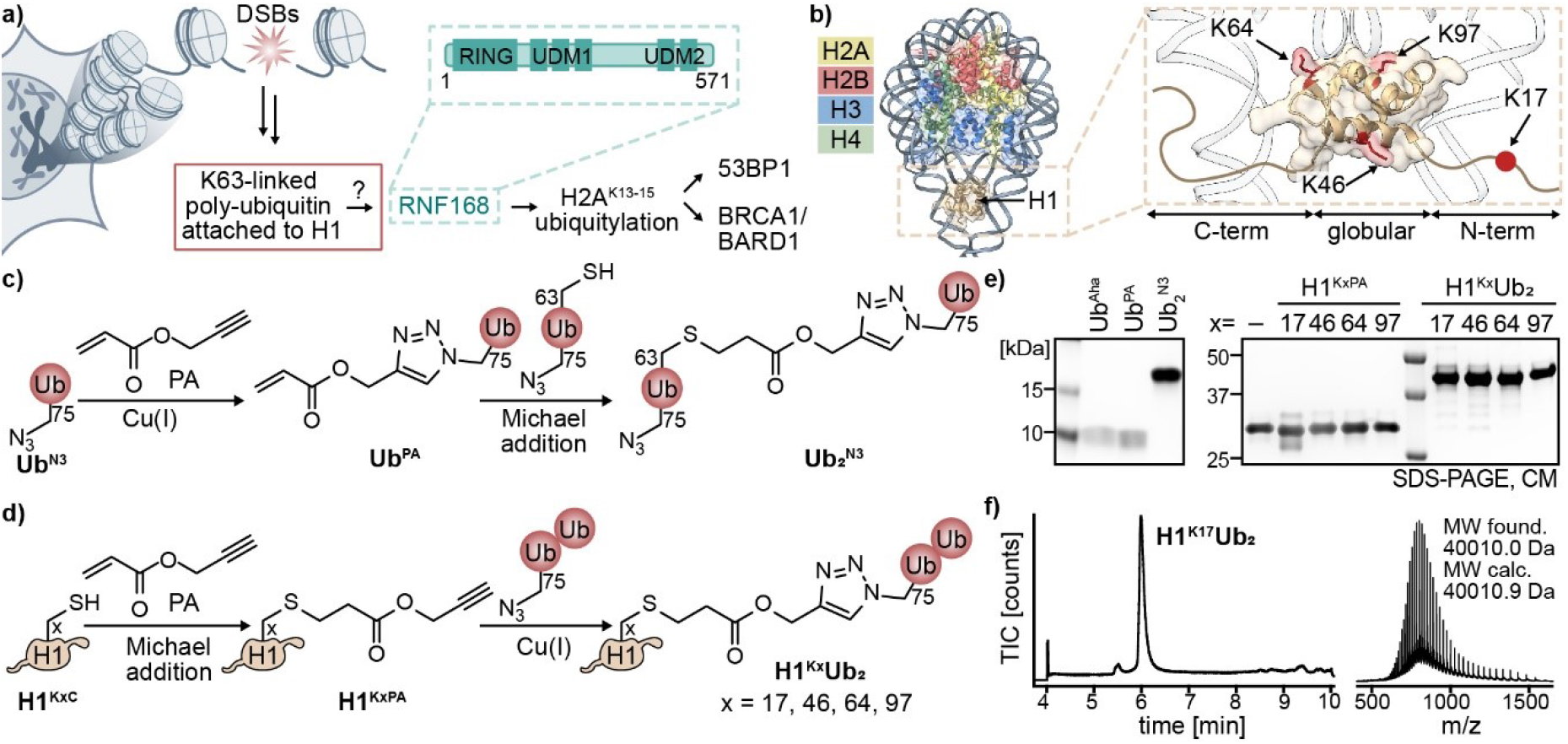
a) The repair of DNA double-strand breaks (DSBs) involves the E3 ubiquitin ligase RNF168 ubiquitylating histone H2A at lysines 13/15, which is recognized by the key DSB repair players p53 binding protein 1 (53BP1) and the heterodimer breast cancer type 1 susceptibility protein (BRCA1)/BRCA1 associated RING domain 1 (BARD1). Recruitment of RNF168 was shown to be facilitated by K63-linked poly-ubiquitin chains, probably attached to H1, yet interactions on a molecular level have been insufficiently addressed. Domain architecture of RNF168 (cyan) indicates the catalytically active RING domain, and two ubiquitin-dependent DSB recruitment modules (UDM 1 and 2). b) Cryogenic electron microscopy (cryo-EM) structure of the chromatosome (PDB:8H0V ^[32]^) with color-coded core histones and H1 positioned at the nucleosomal dyad. Zoom: H1 shows a tripartite structure comprising the globular domain and the intrinsically unstructured N- and C-terminal regions. H1 lysine residues, which are known DNA repair ubiquitylation hotspots^[24]^, are highlighted in red. c) Synthetic strategy to prepare triazole-linked di-ubiquitin (Ub_2_^N3^) using a combination of copper(I)-catalyzed azide-alkyne cycloaddition (CuAAC) and Michael addition. d) Synthetic strategy to attach Ub_2_^N3^ to specific alkyne-modified DNA repair ubiquitylation positions on H1. e) Sodium dodecyl sulfate polyacrylamide gel electrophoresis (SDS-PAGE) analysis of purified ubiquitin variants, H1^Kx^PA, and H1^Kx^Ub_2_ variant. (CM: Coomassie) f) LC-ESI-MS analysis of H1^K17^Ub_2_.(MW: molecular weight; TIC: total ion count)

Chemical methods enable the site-specific installation of ubiquitin moieties on proteins, including histones, and have significantly advanced the mechanistic understanding of chromatin regulation.^[25–27]^ However, most efforts were focused on core histones (**Figure 1b**). H1 family proteins are not part of the nucleosome core particle but bind DNA at the nucleosomal dyad and are critical in regulating chromatin packing (**Figure 1b**).^[28]^ Due to their long, unstructured, and degradation-prone C-terminal domains, H1 subtypes are challenging to handle and chemically modify. Thus, they have not received comparable attention in terms of method development and mechanistic investigations.^[29]^

Here, building on previous approaches to modify H1,^[30,31]^ we used Cu(I)-catalyzed azide-alkyne cycloaddition (CuAAC) to generate a set of specifically K63-linked di-ubiquitylated H1 variants.

Using these molecules *in vitro* and in cells we showed that, indeed, RNF168 activity is stimulated by their presence in a chromatin context, dependent on the modified lysine residue in H1, and that they result in efficient recruitment of RNF168 to chromatin.

## Results and Discussion

### Synthesis of Site-Specifically Di-ubiquitylated H1

The sequence of H1 consists of almost 30% lysines, but only a handful have been found to be poly-ubiquitylated in response to DNA damage.^[24]^ Enzymatic H1 ubiquitylation is promiscuous towards both lysine sites and chain length.^[18]^ Here, we thus used a synthetic approach to couple defined K63-linked di-ubiquitin units, i.e. the shortest ubiquitin-chain recognized by RNF168, site-specifically coupled to four different lysines on H1 that were previously identified as DNA repair-associated ubiquitylation hotspots (K17, K46, K64, and K97, **Figure 1c-f**).^[33]^ Leveraging CuAAC-based ubiquitin polymerization methods,^[34,35]^ we developed a semi-synthetic approach to obtain defined K63-linked di-ubiquitin, connected via a triazole linker. Importantly, these molecules contained an azide at their C-terminus, allowing their subsequent site-specific transfer onto H1.

As a first step, we used selective pressure incorporation (SPI, Fig S1a)^[36]^ to generate a ubiquitin variant bearing the unnatural amino acid azidohomoalanine (Aha) at position 75 in the ubiquitin sequence (Ub^N3^, Fig S1b). We then reacted Ub^Aha^ with propargyl acrylate (PA) via CuAAC to form Ub^PA^ (Fig S1c). In a second step, a ubiquitin variant possessing both a C-terminal Aha residue and a lysine-to-cysteine mutation at position 63 (Ub^N3 K63C^, Fig S1d) was coupled to Ub^PA^ via Michael addition. The reaction yielded the desired K63-linked di-ubiquitin in milligram quantities in isolated yields of up to 45% and bearing a single C-terminal azide for subsequent attachment to H1 (Ub_2_^N3^, **Figure 1c**, Fig S2).

Next, we generated a set of H1 mutants carrying a lysine-to-cysteine mutation at the previously identified ubiquitylation hotspot positions (H1^KxC^, x = 17, 46, 64, 97).^[24]^ The purified H1^KxC^ variants were derivatized with PA via Michael addition to install an alkyne functionality site-specifically at the ubiquitylation sites (H1^KxPA^, Fig S3a-c). CuAAC-based coupling of Ub_2_^N3^ to H1^KxPA^ resulted in the four site-specifically di-ubiquitylated H1 variants (H1^Kx^Ub_2_, **Figure 1d**). Following size exclusion chromatography (SEC), the set of H1^Kx^Ub_2_ variants was obtained in isolated yields ranging from 12% to 38%, primarily depending on the degree of conversion during CuAAC, and in high purity (**Figure 1e,f**, Fig S4).

### RNF168 binds to triazole-linked Ub_2_^N3^

The resulting triazole linker connecting H1 and Ub_2_ is a functional mimic of the native isopeptide bond.^[37–41]^ Typically, rather than directly binding to the linker, ubiquitin chains are recognized by the relative positioning of the ubiquitin moieties towards each other,^[42]^ which is retained in the triazole-linked analogs.^[39–41]^ In the case of RNF168, the crystal structure of the UDM1 domain bound to isopeptide-linked K63-Ub_2_[43] revealed an extended interface of direct contacts between UDM1 and both ubiquitin moieties, but no electron density for the linker itself, suggesting structural flexibility (**Figure 2a**). Still, the differences in chemical nature of the triazole linker and its increased length by three atoms (Fig S3d) may potentially influence the ubiquitin-binding of RNF168. Here, to ensure Ub_2_N3 recognition by RNF168, we examined their interaction experimentally. We used a truncated RNF168 version comprising the RING and UDM1 domains, which has been shown to maintain similar ubiquitylation activity compared to full-length RNF168 *in vitro* (RNF168^1-189^, **Figure 2a**).^[7]^ In pull-down assays, RNF168^1-189^ interacted with both native and triazole-linked K63-linked Ub_2_, indicating that Ub_2_N3 successfully reproduced the native behavior (**Figure 2b**). In contrast, native K48-linked Ub_2_ bound to RNF168^1-189^ with less affinity, demonstrating the specificity of the interaction for the K63 linkage.^[44]^ Together, these results show that the synthetic H1^Kx^Ub_2_ constructs can be used to probe RNF168 regulation.

**Figure 2.**
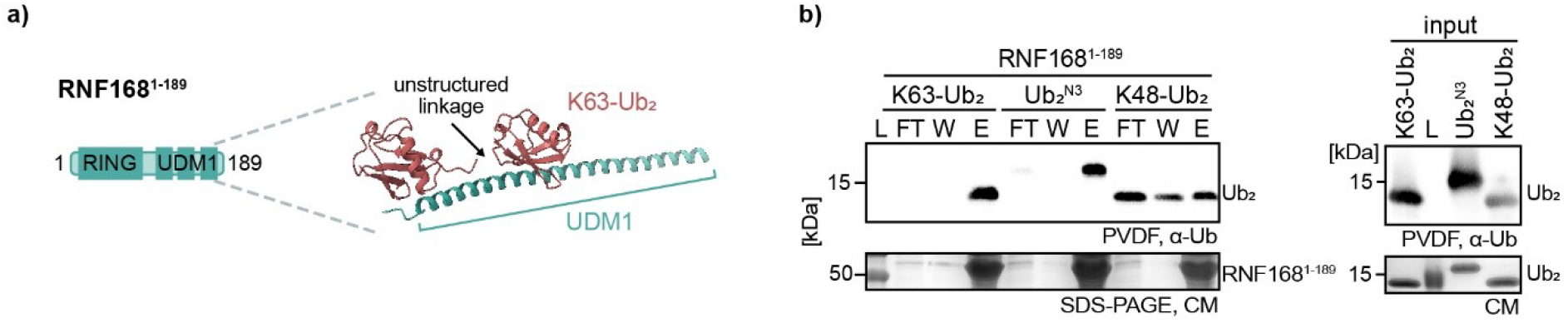
a) Architecture of RNF168^1-189^ showing the RING and UDM1 domains. Close-up shows the crystal structure K63-linked Ub_2_ bound to UDM1 (PDB:5XIS^[43]^). b) Pulldown results: His-tagged RNF168^1-189^ was bound to beads, exposed to Ub_2_^N3^ and native isopeptide-linked K63- and K48-Ub_2_, flowthrough (FT) and wash (W) fractions were collected and the RNF168^1-189^ bound to di-ubiquitin variants was eluted (E) from beads in 200 mM imidazole. Left: Analysis of samples via Western blot (α-Ub, top) and Coomassie blue (CM) staining (bottom). Ub_2_^N3^ showed a slightly higher molecular weight owing to a thrombin cleavage scar. Right: Input for the pull-down assay showing the different di-ubiquitin constructs. (L: ladder)

### Crosstalk of H1 and H2A ubiquitylation via RNF168

Having a set of functional H1^Kx^Ub_2_ constructs in hand, we then reconstituted ‘designer’ chromatin fibers (Fig S5a-b) containing either unmodified H1 or each of the di-ubiquitylated H1 variants (**Figure 3a**). Upon addition of Mg^2+^, the chromatin fibers reversibly condensed into large assemblies that were purified via centrifugation and resuspended in fresh buffer. Throughout this process, both unmodified H1 and the H1^Kx^Ub_2_ variants bound tightly to chromatin fibers, independent of the ubiquitylation site (**Figure 2c**, Fig S5c-d).

**Figure 3.**
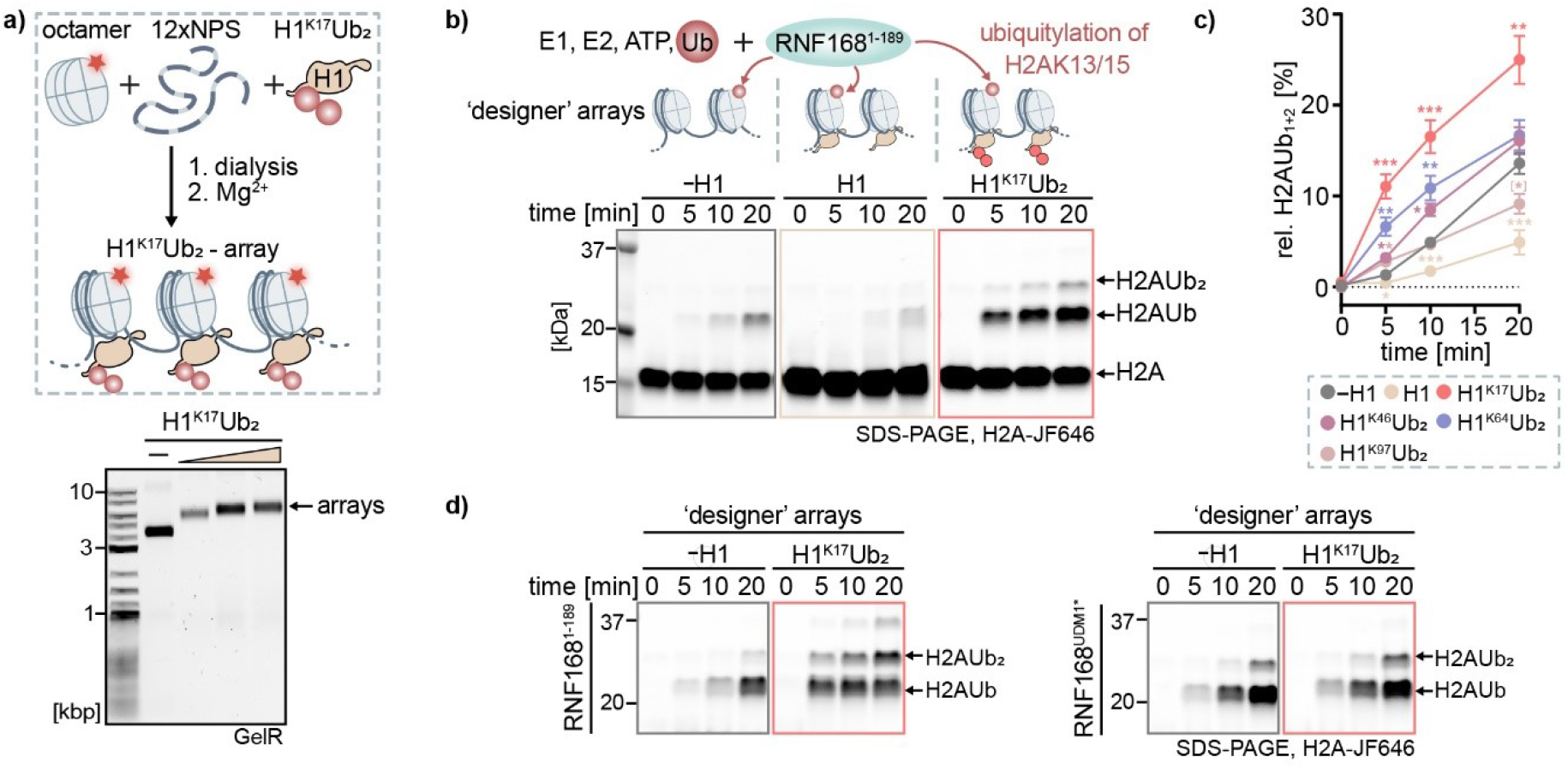
a) Incorporation of H1^K17^Ub_2_ into ‘designer’ nucleosome arrays analyzed by native agarose gel electrophoresis. (GelR: DNA stain) b) SDS-PAGE analysis of in vitro ubiquitylation by RNF168^1-189^ of ‘designer’ nucleosome arrays via in-gel fluorescence of H2A labeled with Janelia Fluor 646 (JF646). For data of other H1^Kx^Ub_2_ variants, see Fig. S6. c) Quantification of in-gel fluorescence intensities corresponding to the sum of the H2AUb and H2AUb_2_ bands normalized to H2A levels at t = 0 min. Data represent mean and standard error of means (SEM) of at least four independent reactions. Significance is indicated for p < 0.05 in relation to the same time point as unmodified chromatin arrays (* = p < 0.05, ** = p < 0.01, *** = p < 0.001). d) SDS-PAGE analysis of in vitro ubiquitylation of H2A in unmodified (-H1) or H1^K17^Ub_2_ containing nucleosome arrays by RNF168^1-189^ (top) or the ubiquitin-binding deficient mutant RNF168^UDM1*^ (bottom). For quantification of the data, see Fig. S7b.

With this set of chromatin fibers, we could test if the presence of one or all variants of H1^Kx^Ub_2_ would influence RNF168 ubiquitylation activity towards H2AK13/15. We thus performed time-resolved ubiquitylation assays with RNF168^1-189^ as E3 ligase, in the presence of free ubiquitin, E1, and E2 enzymes (human UBA1 and UbcH5c, respectively) as well as ATP (**Figure 3b**). Compared to chromatin fibers without H1 (-H1), the presence of unmodified H1 reduced the activity of RNF168 (**Figure 3b,c**). This observation is in agreement with H1 compacting chromatin arrays,^[45]^ rendering H2A less accessible for ubiquitylation. In contrast, the presence of all di-ubiquitylated H1 variants stimulated RNF168 activity, albeit to various degrees (**Figure 3a,b**, Fig S6). A previous study found that mono-ubiquitylated H1 opens chromatin fiber structure,^[31]^ which may result in increased accessibility for RNF168. However, this effect alone cannot explain RNF168 activation by di-ubiquitylated H1 variants compared to unmodified chromatin fibers. Instead, the degree of RNF168 activation varied depending on the Ub_2_N3 attachment site on H1, showing the strongest effect when Ub_2_N3 was placed at position K17 (**Figure 3b,c**). In contrast to the other Ub_2_N3 attachment sites in the globular domain, K17 is positioned on the unstructured N-terminal tail of H1 (**Figure 1b**). This might provide greater flexibility in positioning ubiquitin chains relative to the nucleosome, rendering them more accessible to ubiquitin readers like RNF168. Moreover, the H1^K17^Ub_2_-dependent activation effect was not observed using an RNF168 ubiquitin-binding deficient mutant^[19]^ (RNF168^UDM1*^, **Figure 3c**, Fig S7a-b). Interestingly, we also found that during the ubiquitylation assays, H1^K17^Ub_2_ was poly-ubiquitylated by RNF168^1-189^ but not by RNF168^UDM1*^ (Fig S7c). Considering that neither RNF168 variants ubiquitylated unmodified H1 (Fig S7c), RNF168^1-189^ likely elongated the preinstalled triazole-linked di-ubiquitin on H1. These results indicate that RNF168^1-189^ facilitates its own recruitment, suggesting a positive feedback mechanism. In conclusion, we could show that the binding of RNF168^1-189^ to H1^Kx^Ub_2_ variants, especially at the K17 site, drives the observed increase in H2A ubiquitylation by RNF168.

### H1^K17^Ub_2_ dictates subnuclear localization of RNF168

As a next step, we investigated whether K63-linked di-ubiquitylated H1 is also able to recruit RNF168 to chromatin within the cellular environment. As the cellular context precludes selective control over PTMs, we incorporated the previously synthesized, non-hydrolyzable H1^K17^Ub_2_ into live U-2 OS cells via bead-loading (**Figure 4a**, Fig S8a). Bead-loading offers a straightforward way to deliver proteins (and other macromolecules) into live cells by temporarily disrupting the cell membrane.^[46–48]^ Proteins tolerate this process well, as shown for fluorescently labeled tubulin, which remained functional and was incorporated readily into the cytoskeleton (**Figure 4b**). When bead-loading fluorescently labeled H1 and H1^K17^Ub_2_, the proteins immediately entered the nucleus. There, they displayed a nuclear distribution compatible with random chromatin incorporation, as well as a tendency to accumulate in the nucleolus (**Figure 4b**).

**Figure 4.**
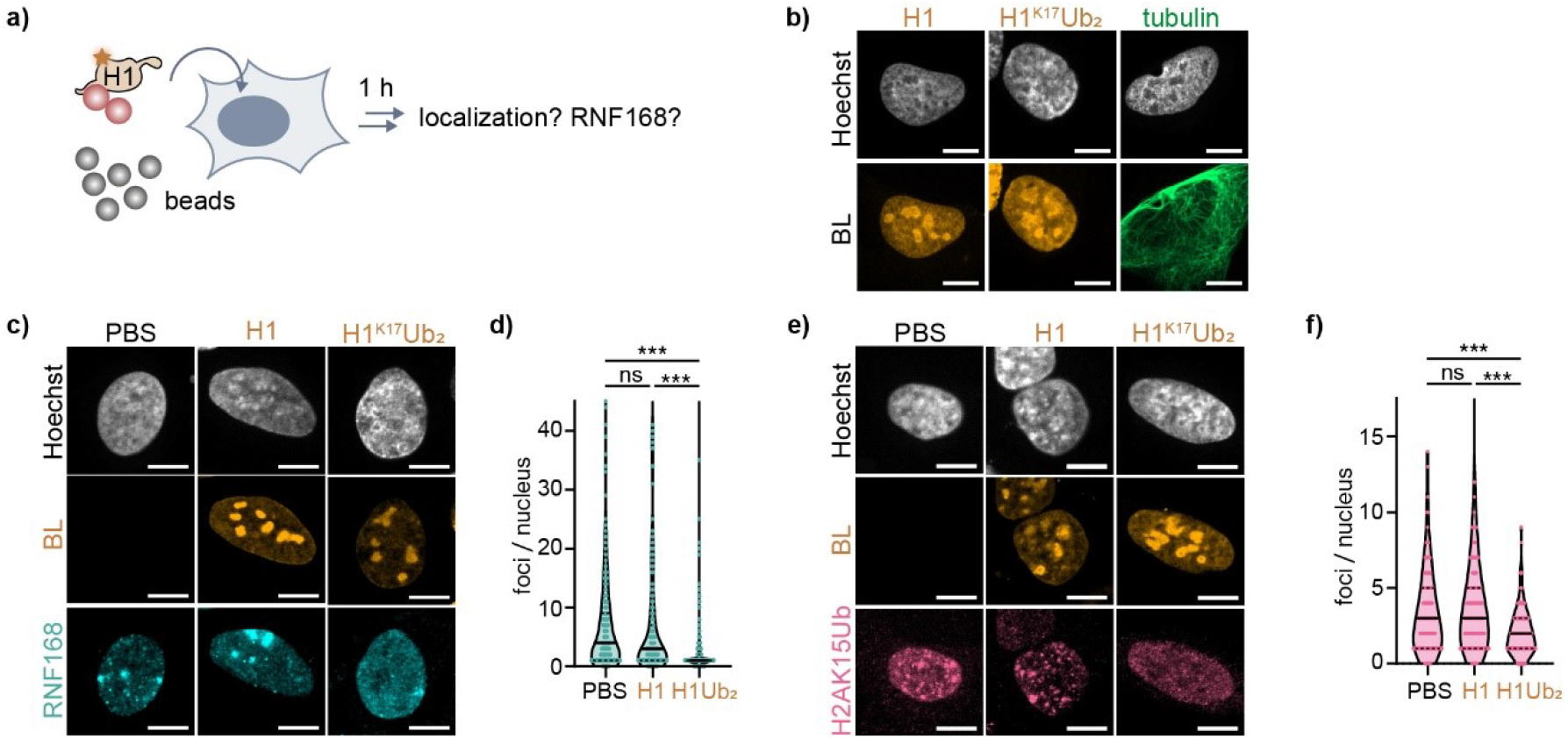
a) Bead-loading (BL) allows the incorporation of macromolecules, including recombinant, PTM-modified proteins such as H1^K17^Ub_2_, into cells. b) Spinning-disk confocal images of fixed U-2 OS cells bead-loaded with fluorescently labeled H1, H1^K17^Ub_2_ (orange), or tubulin (green). c) RNF168 (cyan) was visualized via immunofluorescence (α-FLAG) after BL. d) Quantification of RNF168 foci from c) with data representing four independent replicates with n > 180 nuclei analyzed per condition. e) H2AK15Ub (red) was visualized via immunofluorescence (α-H2AK15Ub). f) Quantification of H2AK15Ub foci per nucleus from e) with data representing two independent replicates with n > 110 nuclei analyzed per condition. (* = p < 0.05, ** = p < 0.01, *** = p < 0.001, scale bar 10 μm)

We then investigated how the presence of ubiquitylated H1 influenced subnuclear RNF168 distribution. We thus used cells expressing FLAG-tagged RNF168 under an inducible promoter (Fig S8b),^[19,20]^ which allowed its detection via an anti-FLAG antibody and immunofluorescence imaging. In cells not exposed to recombinant H1 variants, RNF168 forms distinct foci (**Figure 4c,d**, Fig S8c) associated with DNA replication fork progression via H2A ubiquitylation,^[20]^ as well as sites of endogenous DNA damage. With increasing amounts of randomly deposited H1^K17^Ub_2_, but not H1, FLAG-RNF168 was found to be distributed throughout the nucleus, following H1^K17^Ub_2_ localization (**Figure 4c,d**, Fig S8c). In agreement, the number of distinct H2AK15Ub foci formation also decreased under those conditions and the modification exhibited a more isotropic distribution (**Figure 4e,f**).

To perform the experiments under DNA damage conditions, we treated the cells with the DSB-inducing natural product neocarzinostatin (NCS), which stimulated DNA repair foci formation (Fig S9a). The number of foci containing RNF168 was still suppressed by the random distribution of H1^K17^Ub_2_, but not by H1, alongside H2AK15Ub (Fig S9a,b). Moreover, the recruitment of the DNA damage-associated effector protein 53BP1, which binds to H2AK15Ub within repair foci, was impaired by the introduction of H1^K17^Ub_2_ but not by unmodified H1 (Fig S9c). To further validate that RNF168 recruitment depended on K63-linked ubiquitin binding, we expressed a ubiquitin-binding-deficient RNF168 mutant^[20]^ (RNF168-UBD*). This mutant showed diffuse nuclear localization similar to wild-type RNF168 in the presence of H1^K17^Ub_2_ (Fig S10a,b). Together, these observations indicate that K63-linked ubiquitin installed on H1 directs RNF168 recruitment to chromatin in cells. Accordingly, random chromatin incorporation of H1^K17^Ub_2_ resulted in RNF168 delocalization, its depletion from DNA damage foci, and the inhibition of DNA damage signaling.

## Conclusion

In summary, we expanded the CuAAC-based ubiquitylation toolbox by establishing the generation of defined K63-linked Ub_2_^N3^ in milligram quantities. By attaching Ub_2_^N3^ to four DNA repair-relevant positions on H1, we showed that the different H1^Kx^Ub_2_ variants could be incorporated into chromatin fibers and that they stimulated H2A ubiquitylation by RNF168 in a position-dependent manner. The attachment of Ub_2_ to K17, located on the unstructured N-terminal tail of H1, triggered the strongest RNF168 activation effect, indicating enhanced accessibility for ubiquitin recognition by RNF168. In addition, we showed H1^K17^Ub_2_-dependent chromatin RNF168 recruitment in cells. Recent cryo-EM studies^[16,49]^ show that RNF168 undergoes multiple binding conformations while placing H2AK15Ub, yet large parts of RNF168 (particularly UDM1) remained unresolved, presumably due to molecular flexibility. Thus, we propose that RNF168 UDM1 binding to poly-ubiquitylated H1 guides and stabilizes the initial recruitment step. In conclusion, our results strengthen the hypothesis that direct binding of poly-ubiquitylated H1 by the RNF168 UDM1 domain is a key driving force underlying RNF168 recruitment to DNA damage sites.

## Supporting information

Supporting information is available online.

## Acknowledgments

The authors acknowledge funding from the European Research Council (ERC) (ERC-CoG724022) and EPFL (Ecole Polytechnique Fédérale de Lausanne). We thank Prof. Andreas Marx, Prof. Titia Sixma, and Prof. Michael Rosen for providing plasmids, Dr. Iuliia Boichenko for the preparation of H1.2 mutants, and Prof. Lorenza Penengo for the U-2 OS Flp-In cell lines and feedback on the project. We also acknowledge Olivier Burri (BIOP; EPFL) for help in writing Fiji macros to analyze microscopy data, and for Dr. Carlos Moreno-Yruela for comments on the manuscript.

