## Supporting information for "Chemically di-ubiquitylated H1 variants reveal position-dependent stimulation of the DNA repair protein RNF168"

### Experimental procedures

#### Expression and purification of $Ub^{N3}$, and $Ub^{K63C\ N3}$

The pGEX2TK_GST-Thr-UbG75M and pGEX2TK_GST-Thr-UbK63CG75M plasmids were kindly provided by Andreas Marx (University of Konstanz). The published expression and purification procedures [1,2] were followed, with some adjustments. In brief, plasmids were transformed into methionine (Met) auxotrophic *E.coli* B834 (DE3) bacteria (New England Biolabs, NEB). A pre-culture grown overnight at 37°C in lysogeny broth (LB) supplemented with 100 µg/mL ampicillin was diluted in New Minimal Medium (NMM) containing 0.04 mM methionine (Met, Sigma) and 100 µg/mL ampicillin. Cells were cultured at 37°C until the OD600 reached approximately 1.3, harvested (5000 x g, 20 min, 4°C), resuspended in fresh NMM supplemented with 0.5 mM L-azidohomoalanine (Aha, Roth) and 100 µg/mL ampicillin, and incubated at 37°C for 30 min. Gene expression was induced by adding 1 mM IPTG, and the cells were cultured at 25°C overnight. Cells were harvested via centrifugation (3000 x g, 30 min, 4°C), resuspended in lysis buffer (1% Triton-X 100 in phosphate-buffered saline (PBS) pH 7.4), and lysed via sonication (3x 30 s on, 30 s off, amplitude 20%). Lysates were cleared (50000 x g, 30 min, 4°C), and GST-fusion ubiquitin variants were loaded onto glutathione sepharose beads (GE Healthcare) for 6 h at 4°C. The suspension was transferred into an empty column equipped with a frit, and beads were washed twice with PBS pH 7.4. Ubiquitin mutants $Ub^{N3}$ and $Ub^{K63C\ N3}$ were cleaved on resin by adding 40 U of thrombin (Calbiochem) and incubated at room temperature overnight. The proteins were eluted in PBS pH 7.4 and analyzed by 15% sodium dodecyl sulfate (SDS) polyacrylamide gel electrophoresis (PAGE). Pure fractions were combined, and analyzed by analytical reversed-phase high-performance liquid chromatography (RP-HPLC) and high-resolution electrospray ionization mass spectrometry (HR-ESI-

MS, XEVO G2-XS QTOF) prior to UV-VIS quantification. $Ub^{N3}$ and $Ub^{K63C\ N3}$ were stored as aliquots at −80°C.

**Addition of propargyl acrylate (PA) to $Ub^{N3}$**

$Ub^{N3}$ (150 µM) was mixed with 30 eq. PA (Sigma-Aldrich), which was pre-diluted in acetonitrile (ACN), in click reaction buffer (20 mM Tris-HCl pH 7.5, 3 mM tris(3-hydroxypropyltriazolylmethyl)amin (THPTA), 1.5 mM $CuSO_4$, 15 mM sodium L-ascorbate). The final concentration of ACN was 10%. The reaction was incubated on ice and monitored by LC-ESI-MS. Upon quantitative conversion, the reaction was quenched by the addition of 50 mM ethylenediaminetetraacetic acid (EDTA). Excessive PA and other reaction components were removed by dialysis (50% methanol in 20 mM Tris-HCl, 150 mM NaCl, 1 mM EDTA) step-wise, reducing methanol to 0% until PA was removed completely (usually 1 d to 2 d). Finally, $Ub^{PA}$ was UV-VIS quantified, aliquoted, and stored at −80°C.

**Preparation of $Ub_2^{N3}$**

$Ub^{PA}$ (150 µM) and $Ub^{K63C\ N3}$ (1.1 eq.) were mixed in 20 mM Tris base, and incubated overnight at 22°C. The reaction was centrifuged (21’000 x g, 10 min, 4°C) and purified via size exclusion chromatography (SEC) using a Superdex 75 10/300 GL column (GE Healthcare) in SEC buffer (20 mM Tris-HCl pH 7.5, 150 mM NaCl). Fractions were analyzed by SDS-PAGE, and pure fractions were collected, quantified by UV-VIS spectroscopy, and analyzed by analytical HPLC and HR-ESI-MS.

**Expression and purification of H1 and $H1^{KxC}$**

Single point mutations (Lys to Cys) were introduced via Gibson assembly to the plasmid pET11a_H1.2WT[3] to obtain pET11a_H1.2$^{KxC}$ (x = 17, 46, 64, 97). H1 variants were expressed in BL21(DE3) E.coli (Agilent) by addition of 1 mM isopropyl β-D-1-thiogalactopyranoside (IPTG) for 3 h at 30°C. Cells were harvested (3000 x g, 30 min, 4°C), resuspended in pre-cooled lysis buffer (20 mM Tris-HCl pH 7.5, 150 mM NaCl, 0.2 mM 4-(2-aminoethyl)benzenesulfonyl fluoride hydrochloride (AEBSF), 1 mM dithiothreitol (DTT), 0.1 % Triton X-100, 1 x protease inhibitor cocktail (Roche)), and lysed via sonication (1 s on, 2 s off, 2 min, amplitude 35 %). RNase A (Qiagen) and DNase (Sigma) were added to a final concentration of 1 µg/mL, and lysates were cleared via centrifugation (50’000 x g, 30 min, 4°C). The supernatant was purified by ion exchange chromatography (IEX) using a 5 mL HiTrap SP HP (Cytiva) column and the following buffers: 20 mM Tris-HCl pH 7.5, 1 mM EDTA, 150 (buffer A) and buffer B: similar composition as A but containing 2000 mM NaCl. The protein was purified employing a gradient of 0 % to 50 % B in A over 20 column volumes (CV). The relevant fractions were combined after SDS-PAGE analysis. H1 was further purified by semi-preparative RP-HPLC and the fractions were analyzed by HR-ESI-MS. Pure fractions were combined, lyophilized, and stored as a powder at −80°C. $H1^{KxC}$ variants were prepared similarly to H1. After IEX, the concentration of $H1^{KxC}$ was determined via UV-VIS. 20 µM $H1^{KxC}$ were reacted with 50 eq. propargyl acrylate (PA) in reaction buffer (20 mM Tris-HCl pH 7.5, 150 mM NaCl, 10 % ACN) for 30 min to 60 min at ambient temperature. Upon full conversion monitored by high-resolution liquid chromatography electrospray mass spectrometry (HR-LC-ESI-MS XEVO G2-XS QTOF), $H1^{KxPA}$ variants were purified via semipreparative RP-HPLC. The pure fractions analyzed by HR-ESI-MS were combined, lyophilized, and stored as a powder at −80°C. For quality control, H1 and $H1^{KxPA}$ were analyzed by analytical RP-HPLC and LC-ESI-MS.

**Chemical ubiquitylation of H1 ($H1^{KxC}Ub_2$)**

$H1^{KxPA}$ (30 µM to 50 µM) was mixed with $Ub_2^{N3}$ (1.5 eq.) in click reaction buffer (20 mM Tris-HCl, 20% DMSO, 10 mM THPTA, 5 mM $CuSO_4$). 50 mM Na-ascorbate was added last to start the reaction. Argon was bubbled through the reaction, and the mixture was incubated at 22°C, protected from light for 1 h. The reaction was quenched by the addition of 50 mM EDTA, centrifuged (21’000 x g, 10 min, 4°C), and purified by SEC using a Superdex200Increase 10/300 GL column (Cytvia). Fractions were analyzed by SDS-PAGE, pure fractions were collected, quantified by UV-VIS spectroscopy, analyzed by HR-LC-ESI-MS, and stored at –80°C.

**Expression and purification of $RNF168^{1\text{-}189}$, $RNF168^{UDM1*}$, and $RNF168^{RING}$**

The pETNKI_6xHis-SUMO3-$RNF168^{1\text{-}189}$ plasmid was a gift from the Titia Sixma laboratory (Netherlands Cancer Institute).[4] The plasmid was transformed into Rosetta-gami2 (DE3) (Merck Millipore) cultured in LB supplemented with 1 µM ZnCl2, 50 µg/µL kanamycin and 24 µg/µL chloramphenicol were diluted in LB with similar composition and grown until an OD600 value of approximately 0.6 was reached. Gene expression was induced by adding 0.25 mM IPTG, and cells were cultured at 16°C overnight. Cells were harvested (5000 x g,20 min,4°C), resuspended in pre-cooled lysis buffer (20 mM Tris-HCl pH 7.5, 500 mM NaCl, 10% glycerol, 1 µM ZnCl2, 1 mM DTT, 2 mM PMSF, one protease inhibitor pill (EDTA-free, Roche) per 50 mL), and lysed via sonication (3 min 1 s on, 1 s off, amplitude 40%). Cell lysates were cleared (23’000 x g, 30 min, 4°C), and incubated with ZnNTA beads (Cube-Biotech) for 1 h at 4°C. The suspension was transferred into an empty column equipped with a frit, and beads were washed twice with 10 column volumes (CV) pre-cooled lysis buffer and twice with 10 CV lysis buffer containing 5 mM imidazole. Proteins were eluted with an imidazole gradient (10 mM to 500 mM), and fractions were analyzed by 15% SDS-PAGE. Pure fractions were combined, and the 6xHis-SUMO3 tag was cleaved by adding 20 nM SENP2 (Boston Biochem) while dialyzing against cleavage buffer (50 mM Tris-HCl pH 7.5, 150 mM NaCl, 0.2% NP-40, 1 µM ZnCl2, 1 mM DTT) at 4°C overnight. $RNF168^{1\text{-}189}$ was further purified by reverse ZnNTA affinity and size exclusion chromatography (SEC) using a Superdex 75 10/300 GL (GE Healthcare) column in SEC buffer (50 mM Tris-HCl pH 7.5, 150 mM NaCl, 0.2% NP-40, 1 µM ZnCl2, 5 mM beta-mercaptoethanol). RNF168$^{1\text{-}189}$ was UV-Vis quantified, aliquoted, and stored at −80°C.

**Preparation of nucleosome array DNA containing 12 nucleosome positioning sequences (12xNPS)**

The pWM_12x601_192NRL plasmid was a kind gift from the Michel Rosen laboratory (UT Southwestern).[5] It was transformed in dam$^-$/dcm$^-$ competent E.coli (NEB), and stored as glycerol stocks at −80°C. A pre-culture grown in LB supplemented with 100 µg/mL ampicillin was diluted in LB with similar composition, and cultured at 37°C overnight. Cells were harvested (3000 x g, 30 min, 4°C), and suspended in pre-cooled lysis buffer I (50 mM glucose, 25 mM Tris-HCl pH 7.0, 10 mM EDTA pH 8.0). Subsequently, three volumes of lysis buffer II (0.2 M NaOH, 1% SDS) and three volumes lysis buffer III (4 M KAc, 2 M AcOH) were added. Cell debris was removed by centrifugation (12'000 x g, 15 min, 4°C), supernatant was filtered (Miracloth), and genetic material was precipitated by isopropyl alcohol (iPr). Plasmid DNA was harvested by centrifugation (12'000 x g, 15 min, 4°C), air dried, and re-dissolved in TE 10/50 (10 mM Tris-HCl pH 7.0, 50 mM EDTA pH 8.0). RNA was digested (0.3 mg/mL RNase (Qiagen)) for 8 h at 37°C. Plasmid DNA was separated from RNA fragments via SEC using a Sepharose 6 XK30/50 column (GE Healthcare) in TEK2000 buffer (10 mM Tris-HCl pH 7.0, 50 mM EDTA pH 8.0, 2 M KCl). Fractions were analyzed by 1% agarose gel electrophoresis, combined and iPr-precipitated. Plasmid DNA was re-dissolved in TE 10/0.1 buffer (10 mM Tris-HCl pH 7.0, 0.1 mM EDTA pH 8.0), its concentration was determined via UV-VIS spectroscopy, and stored at −20°C. The 12xNPS nucleosome array DNA insert was released from the vector backbone by EcoRV digestion (440 units (U) /mL, NEB) in CutSmart buffer (NEB) for 4 d at 37°C. The DNA fragment was precipitated with increasing concentrations of polyethylene glycol (PEG), harvested (15'000 x g, 30 min, 4°C), and re-dissolved in TE 10/0.1. The DNA was further purified by chloroform-phenol extraction (Sigma), and precipitation with ethanol and 3 M sodium acetate. Pure 12xNPS was harvested via centrifugation (15'000 x g, 30 min, 4°C), re-dissolved in TE 10/0.1, quantified via UV spectroscopy, and aliquots were stored at −20°C. The purification procedure was monitored by 1% agarose gel electrophoresis.

**Octamer assembly**

Human core histones H2A, H2A$^{N110C}$, H2B, H3$^{C110A,}$ and H4 were expressed and purified as described previously.[2] H2A$^{N110C}$ was used for fluorescent labeling. Purified H2A$^{N110C}$ (0.56 µM) was dissolved in reaction buffer (20 mM Tris-HCl pH 8.5, 150 mM), and tris(2-carboxyethyl)phosphine (TCEP, 2 eq., pH 7.5 - 8.0) was added, and the solution was incubated at 25°C for 30 min. Then, Janelia Fluor

646 maleimide (JF646, 2.5 eq., Tocris) was added. Quantitative conversion was observed after 20 min following the reaction by analytical RP-HPLC and HR-ESI-MS analysis. $H2A^{N110C}$JF646 was purified by semi-preparative RP-HPLC. The purified product was analyzed by analytical RP-HPLC and LC-ESI-MS. Histone octamers were assembled as previously described. Briefly, purified core histones were dissolved in unfolding buffer (6 M guanidinium hydrochloride, 10 mM Tris-HCl pH 7.5, 5 mM DTT). Samples were centrifuged (21'000 x g, 10 min, 4°C), and the concentration was determined using UV-VIS spectroscopy using the following molar extinction coefficients at 280 nm: $\varepsilon_{280}$(H2A) = 4470 $M^{-1}cm^{-1}$, $\varepsilon_{280}$(H2B) = 7450 $M^{-1}cm^{-1}$, $\varepsilon_{280}$($H3^{C110A}$) = 4470 $M^{-1}cm^{-1}$, $\varepsilon_{280}$(H4) = 5960 $M^{-1}cm^{-1}$. H3 (1 eq.) and H4 (1 eq.) were mixed with H2A (1.05 eq.) and H2B (1.05 eq.) to obtain a 1 mg/mL solution, which was dialysed against refolding buffer (2 M NaCl, 10 mM Tris-HCl pH 7.5, 1 mM EDTA, 5 mM DTT) at 4°C overnight. Refolded octamers were purified via SEC using a Superdex 200 10/300 GL column (GE Healthcare) in refolding buffer. Fractions were analyzed by 17% SDS-PAGE, combined, concentrated up to a final concentration of 20 µM, and stored in 50% glycerol at −20°C.

**Nucleosome array reconstitution**

Nucleosome arrays were assembled as published previously.[6] In short, reconstitution at a final concentration of 1 µM per nucleosome was performed by mixing 12xNPS DNA, appropriate equivalents of H2A-JF646 containing octamers, and buffer DNA (0.5 eq.MMTV DNA) in 2 M NaCl. H1 or $H1^{Kx}Ub_2$ variants were added when indicated. Mixtures were gradually dialyzed from TEK2000 (10 mM Tris-HCl pH 7.5, 0.1 mM EDTA, 2000 mM KCl) to TEK10 (10 mM Tris-HCl pH 7.5, 0.1 mM EDTA, 10 mM KCl) using a peristaltic pump. Chromatin concentration was quantified by UV-VIS spectroscopy, and analyzed by native agarose gel electrophoresis. Nucleosome array compaction was induced by the addition of 4 mM $Mg^{2+}$. Arrays were incubated for 20 min at 4°C, centrifuged (21'000 x g, 10 min, 4°C), and gently resuspended in TEK10. After analysis by native agarose gel electrophoresis, arrays were used for ubiquitylation assays.

**RNF168 ubiquitylation assays**

$RNF168^{1-189}$ (0.125 µM), nucleosome arrays (0.25 µM), ubiquitin (50 mM, Boston Biochem), and UbcH5c/UBE2D3 (0.9 µM, Boston Biochem) were mixed in 1x ubiquitylation buffer (30 mM 4-(2-hydroxyethyl)-1-piperazineethanesulfonic acid (HEPES) pH 7.5, 5 mM $MgCl_2$, 0.2 mM DTT, 10 mM sodium citrate, 20 mM ATP, 10 mM creatine phosphate, 0.2 µg/mL creatine phosphokinase). The E1 ubiquitin ligase UBE1 (0.3 µM, Boston Biochem) was added last. The reaction was monitored by withdrawing samples after indicated time intervals, quenched by the addition of 5x SDS sample buffer (25 mM Tris base pH 6.8, 1% SDS, 50% glycerol, 0.005% bromophenol blue, 10% beta-mercaptoethanol), heated at 95°C for 10 min, and analyzed by 15% SDS-PAGE. Band intensities were quantified by ImageJ using the built-in Gel analyzer tool. Data was plotted in GraphPad Prism, and statistical significance was calculated using the two-way ANOVA with Dunnet's multiple comparison test. Results were accepted as significantly different for $p < 0.05$.

**Pull-downs**

Bacterial lysate containing 6xHis-tagged RNF168$^{1\text{-}189}$ from expression cultures was thawed at ambient temperature, sonicated and loaded onto a column containing resin beads functionalized with nitrilotriacetic acid (NTA) and complexed with Zn (ZnNTA) for 1 h at 4°C. The suspension was transferred to an empty column equipped with a frit and washed with 3x 10 CV pre-cooled binding buffer (50 mM Tris-HCl pH 7.5, 150 mM NaCl, 1 µM $ZnCl_2$, 5 mM beta-mercaptoethanol). Immobilized RNF168$^{1\text{-}189}$ was incubated with 400 pmol ubiquitin-dimers (native K48-, K63- and triazole-linked K63-$Ub_2$) for 1 h at 4°C. Subsequently, beads were washed with 2 x 10 CV binding buffer and 2 x 10 CV binding buffer containing 5 mM imidazole. RNF168$^{1\text{-}189}$ was eluted in 200 mM imidazole. Flowthrough, wash, and elution fractions were analyzed by 14% SDS-PAGE and Western Blotting using an α-ubiquitin antibody (α-Ub, 1:000, ab19247 Abcam). GST-tagged RNF168 variants (2.5 µM) were incubated with H1, $Ub_2$ or H1$^{K17}$$Ub_2$ (5 µM) for 30 min at 30°C in binding buffer 20 mM Tris-HCl, 150 mM NaCl, 1 mM $ZnCl_2$, 1 mM DTT, 0.1% NP-40, 0.2 mg/mL bovine serum albumin (BSA)). Subsequently, pre-equilibrated magnetic glutathione (GSH)-agarose beads (Pierce) were added and incubated shaking (500 rpm) at ambient temperature for 1.5 h. Then, beads were washed 2 x with 10 CV binding buffer, and resuspended in milliQ water. Beads were boiled at 95°C for 5 min, and the supernatant was analyzed by 15% SDS-PAGE and Western Blot using the following antibodies: α-Ub, 1:000 (ab19247, Abcam), α-H1, 1:1000 (ab17677, Abcam).

**Cell culture**

All cells were maintained at 37°C and 5 % CO2 in a humidified atmosphere, hereafter referred to as standard culture conditions. U-2 OS cells were cultivated in McCoy 5A Medium (Sigma-Aldrich) containing 10 % fetal bovine serum (FBS), 100 U/mL penicillin, and 100 µg/mL streptomycin. U-2 OS Flp-In TREx FLAG-RNF168 (U-2 OS RNF168) cells were a gift from Lorenza Penengo (University of Zurich)[7] and cultured in Dulbecco's modified Eagle's medium (DMEM, Sigma-Aldrich) containing 10 % tetracycline-free FBS (Gibco) in the presence of 100 U/mL penicillin, 100 µg/mL streptomycin, 5 µg/mL blasticidin, and 5 µg/mL hygromycin. Cells were routinely subcultured twice a week at 70% to 80% confluency. The total number of cells and the proportion of viable cells were determined using a Neubauer chamber, and cells were only used for experiments when the viable fraction of cells was > 90% and showed a constant doubling time. Cells were used for experiments from passages four to twelve.

**Bead loading**

The procedure was performed essentially according to Mann et al.[8] with minor adjustments. Cells were seeded in an 8- or 18-well μ-slide with polymer coverslip (ibiTreat or poly-lysine coating, Ibidi) to obtain 70% to 80% confluency at the day of bead-loading. Non-fluorescent proteins were labeled with JF549-NHS (1.2 eq.) for 30 min at ambient temperature in 100 mM $NaHCO_3$. The medium was removed, and cells were washed with pre-warmed PBS containing 0.1% pluronic F-68 (PBS-F). Proteins were diluted appropriately in PBS-F, and added to the cells. A thin layer of beads was sprinkled onto the cells, and plates were tapped eight times while gently swirling the culture dish. Cells were allowed to recover for 2 min at 37°C. Then, the protein solution was removed by gentle pipetting, and cells were washed once with PBS-F, and twice with medium to remove the beads. Cells were allowed to recover in medium for 30 min to 60 min before fixation. If indicated, cells were treated with 100 ng/mL neocarzinostatin (NCS) for 1 h prior and post bead-loading. For immunofluorescence, cells were washed three times with pre-warmed PBS, and then fixed in 4% para-formaldehyde (PFA) (Electron Microscopy Sciences, EMS) in PBS for 10 min at ambient temperature. Cells were re-washed three times with PBS and permeabilized with 0.2% Triton X-100 in PBS for 5 min, followed by blocking in 5% BSA in PBS for 2 h. Afterwards, cells were washed with 0.1% Triton in PBS, and incubated with primary antibodies in 5% BSA in PBS for 2 h (α-γH2A.X 1:400 (2577S, Cell Signaling Technologies), α-FLAG 1:1000 (F7425, Sigma Aldrich), α-H2AUb 1:200 (MABE1119, Sigma Aldrich), α-FLAG 1:300 (F7425, rabbit, Sigma Aldrich), α-FLAG 1:1000 (F1804, mouse, Sigma Aldrich), α-53BP1 1:300 (612522, BD Biosciences)). Cells were washed three times with 0.1% Triton in PBS, and then incubated with suitable secondary antibodies (1:1000 in 5% BSA in PBS containing 2 μg/μL Hoechst). Cells were imaged on a Nikon Spinning Disk CSU W1 with an SR HP Apo TIRF 100x/1.49 oil objective. Image analysis was performed using a Fiji Macro, which included background subtraction, nuclei detection via Hoechst staining, and foci counting within nuclei borders. Thresholds were kept constant for all experiments. Nuclei were excluded from analysis if (1) cells did not express RNF168 and/or (2) were not bead-loaded, and/or (3) were aberrantly small or large (accepted 120 $\mu m^2$ to 460 $\mu m^2$). Foci data was plotted using GraphPad Prism, and statistical significance was calculated using Kruskal Wallis and Dunn's multiple comparison test. Results were accepted as significantly different for $p < 0.05$.

### Supplementary Figures

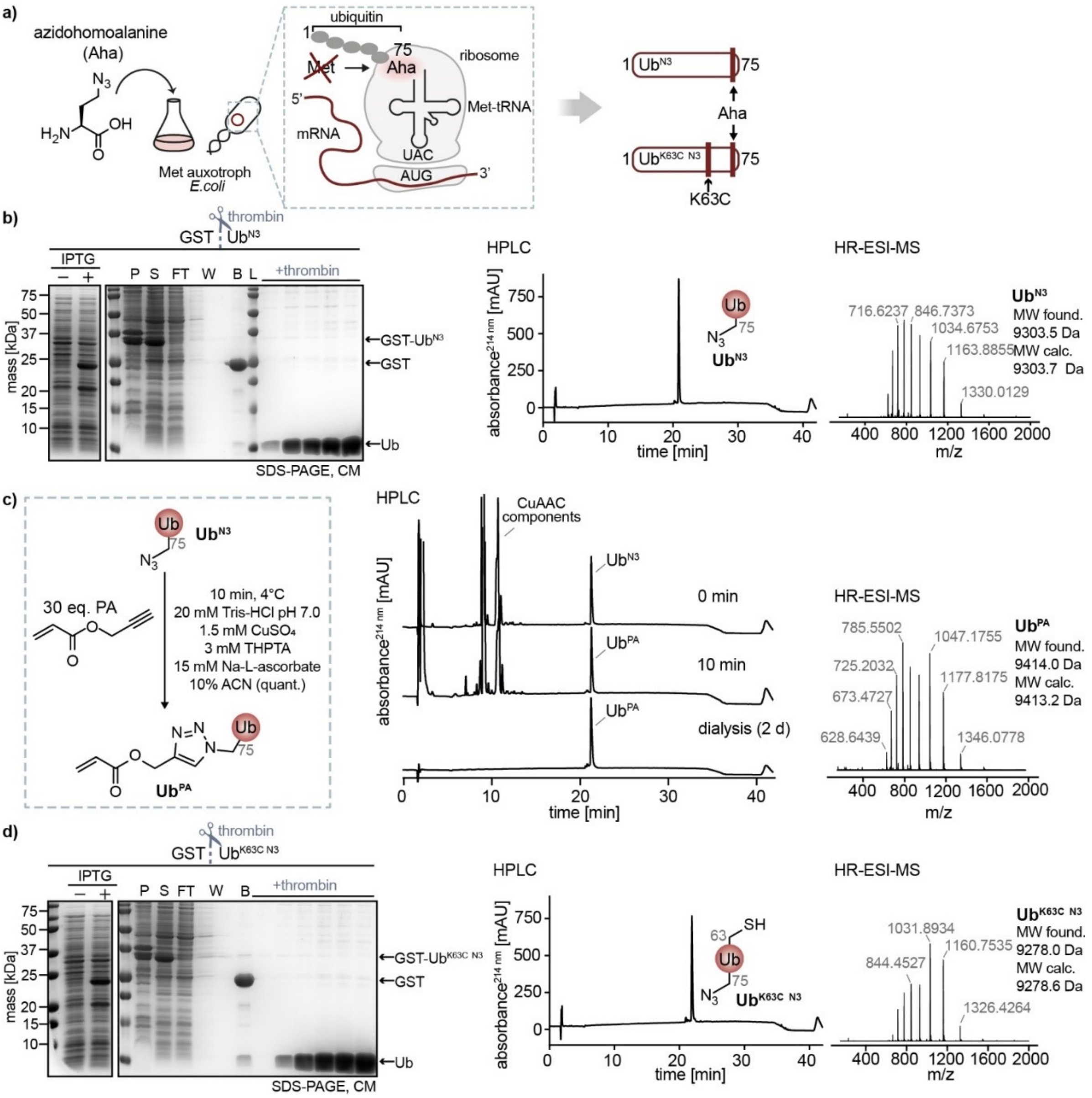


**Figure S1.** a) Illustration showing selective pressure incorporation (SPI) of azidohomoalanine (Aha) into $Ub^{N3}$ and $Ub^{K63C\ N3}$. In methionine-auxotroph bacteria cultured in methionine-free media, Aha is translationally incorporated instead of methionine. The ubiquitin C-terminal glycine 75 was replaced by methionine, and glycine 76 was deleted. $Ub^{K63C\ N3}$ carried a lysine-to-cysteine mutation at position 63. b) Analysis of expression and GST-affinity purification of $Ub^{N3}$ analyzed by SDS-PAGE. Analytical HPLC and HR-ESI-MS analysis of purified $Ub^{N3}$ confirmed quantitative incorporation of Aha. c) Addition of PA to $Ub^{N3}$ via CuAAC to form $Ub^{PA}$. Reaction and dialysis monitoring via analytical HPLC and HR-ESI-MS. d) SDS-PAGE analysis of expression and GST-affinity purification of $Ub^{K63C\ N3}$. Analytical HPLC and HR-ESI-MS analysis of $Ub^{K63C\ N3}$ purified confirmed quantitative incorporation of Aha.

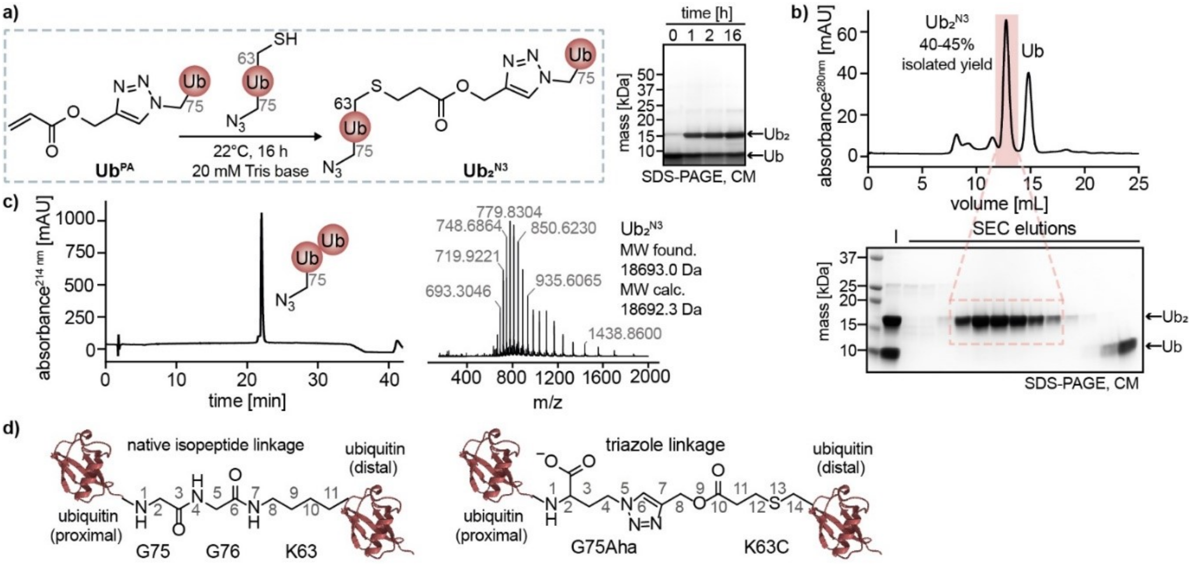


**Figure S2.** a) Schematic overview and SDS-PAGE analysis of Ub$^{PA}$ and Ub$^{K63C\ N3}$ coupling via Michael addition to form Ub$_2^{N3}$. b) Preparative-scale preparation of Ub$_2^{N3}$ purified by size-exclusion purification (SEC) and analyzed by SDS-PAGE. c) Analytical HPLC and HR-ESI-MS analysis of Ub$_2^{N3}$. d) Schematic representation of native isopeptide-linked and triazole-linked Ub$_2$ (PDB:1UBQ[9]).

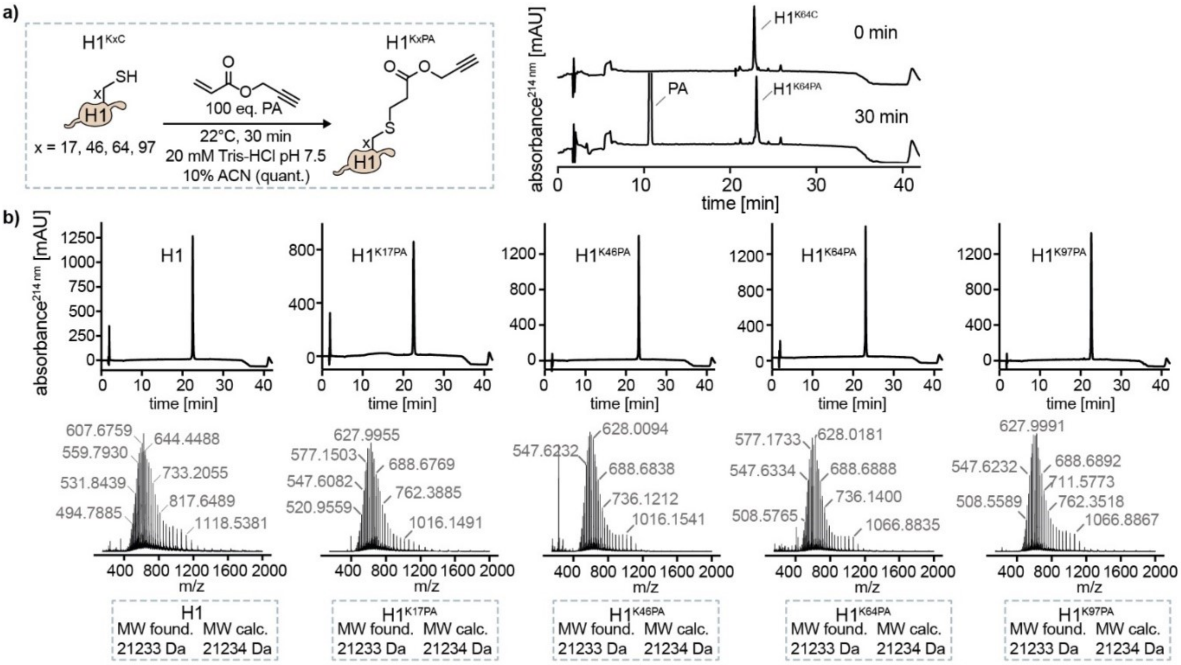


**Figure S3.** a) Schematic overview of the optimized reaction conditions for the Michael Addition of PA to H1$^{KxC}$ (20 μM). b) Representative reaction monitoring of H1$^{K64}$PA generation by analytical HPLC. c) Analytical HPLC and HR-ESI-MS analysis of all H1$^{KxPA}$ variants after preparative HPLC purification.

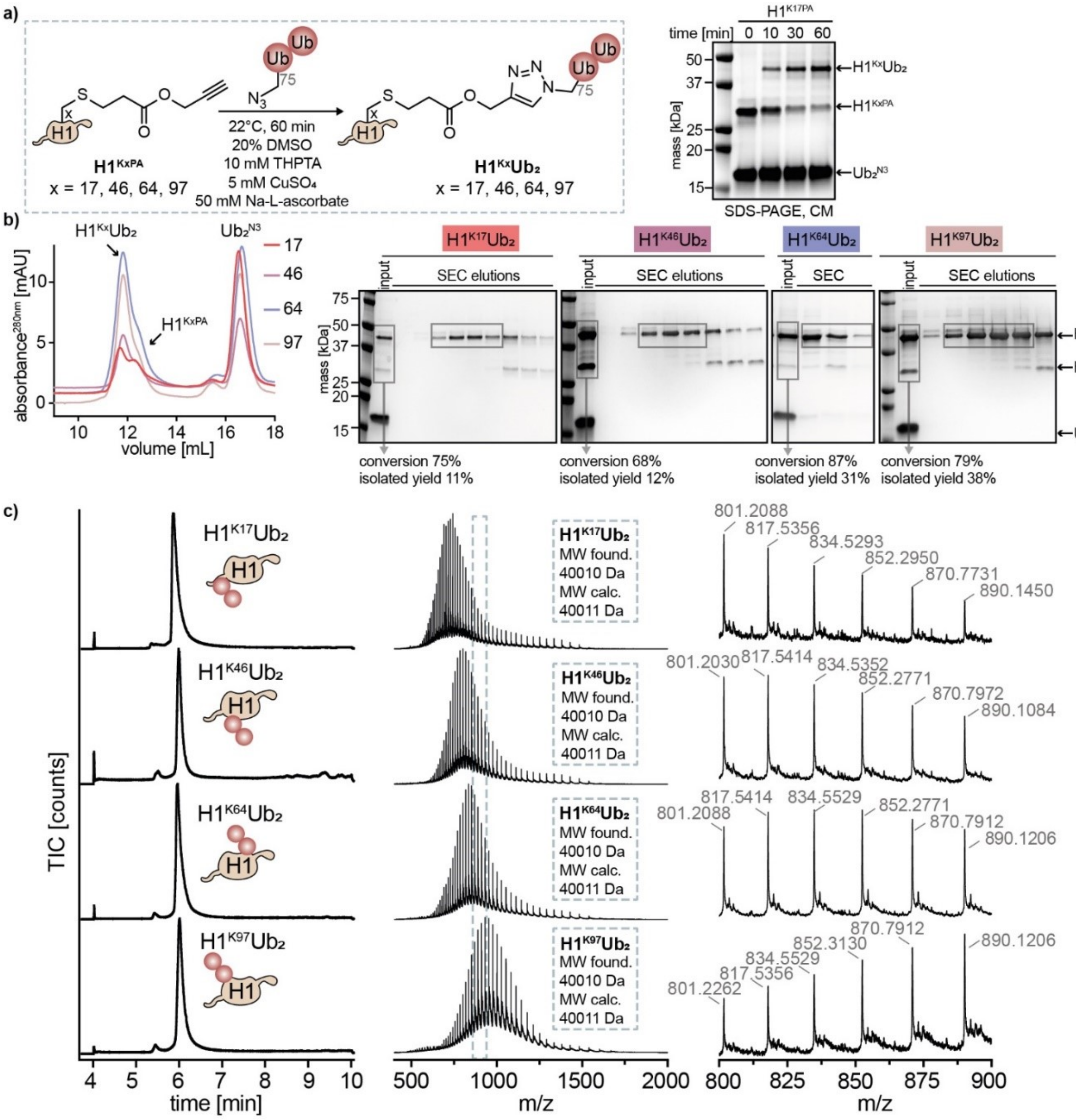


**Figure S4.** a) Schematic overview of CuAAC-mediated coupling of $Ub_2^{N3}$ and $H1^{Kx}PA$. Final optimized reaction conditions are indicated. Representative, time resolved SDS-PAGE analysis of $H1^{K17}PA$ reacting with $Ub_2^{N3}$. b) SEC profiles with corresponding SDS-PAGE analyses of $H1^{Kx}Ub_2$ variants purification. Conversion rates and isolated yields are indicated. c) HR-LC-ESI-ES analysis of $H1^{Kx}Ub_2$ variants: total ion count (TIC) chromatograms (left), m/z distribution (middle), and a zoom-in view of the m/z distribution from 800 m/z to 900 m/z (right).

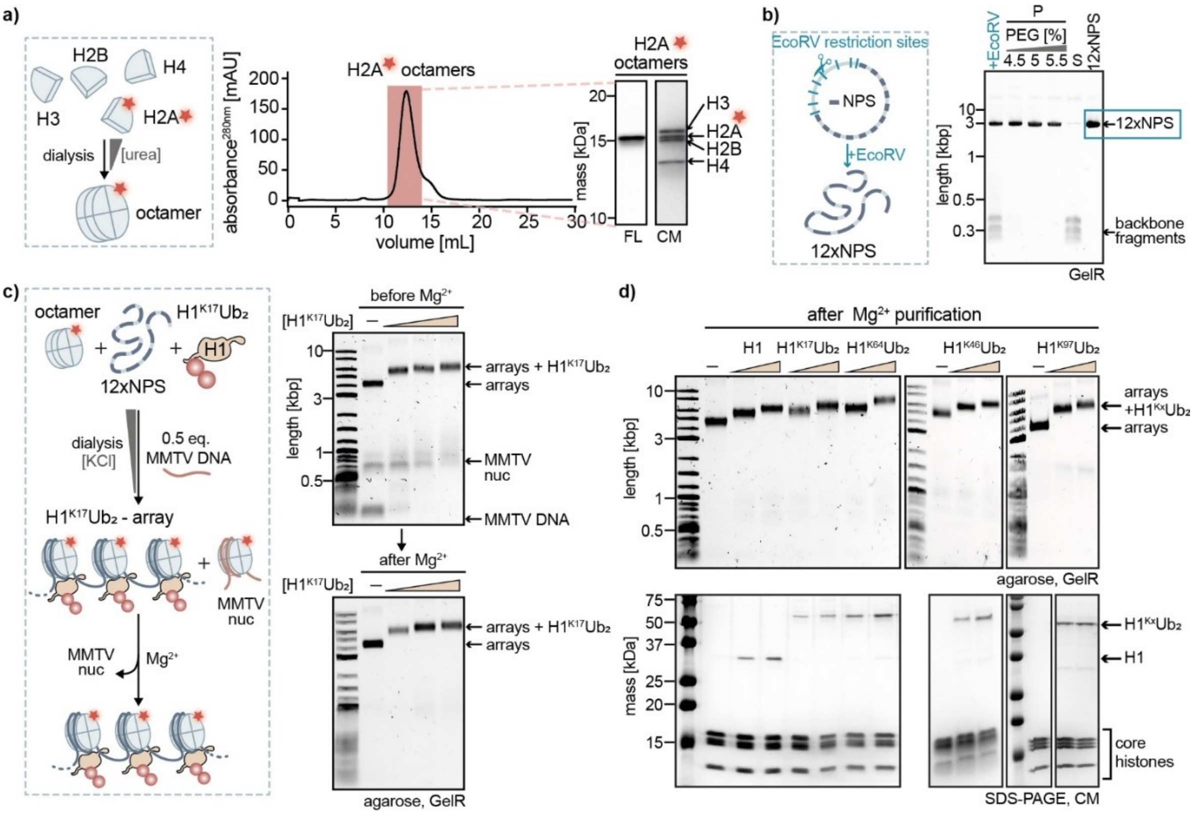


**Figure S5.** a) Octamer assembly with fluorescently labeled H2A (H2A-JF646) purified by SEC and analyzed by SDS-PAGE. b) After EcoRV restriction digestion, PEG purification allowed the removal of backbone fragments resulting in pure 12xNPS DNA, as shown by agarose gel electrophoresis. c) Schematic overview of $H1^{K17}Ub_2$-nucleosome array reconstitution, subsequent $Mg^{2+}$-induced pelleting, and resuspension to remove MMTV nucleosomes (MMTV nuc) monitored by native agarose gel electrophoresis. d) Native agarose gel electrophoresis (top) and SDS-PAGE analysis (bottom) of H1 and $H1^{Kx}Ub_2$-nucleosome arrays after $Mg^{2+}$-purification.

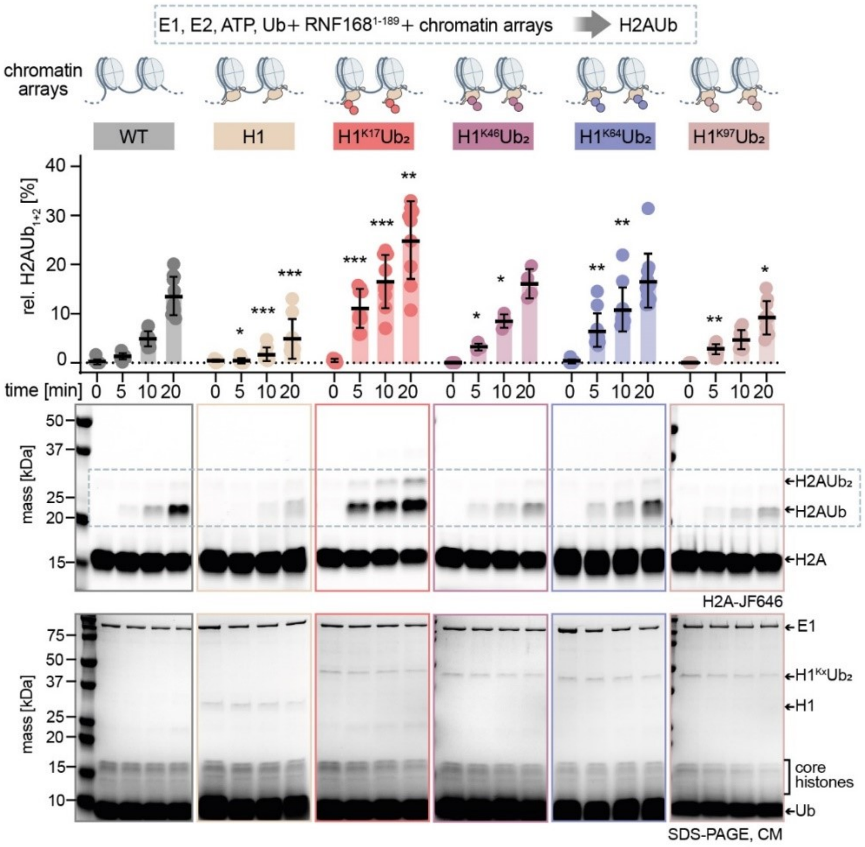


**Figure S6.** Additional data to Figure 3b and c: Quantification of in-gel fluorescence intensities (H2A-JF646) corresponding to the sum of the H2AUb and $H2AUb_2$ bands normalized to H2A levels at t = 0 min. Data shows the mean and standard deviation of at least four independent reactions with individual data points indicated. Significance is indicated for $p < 0.05$ in relation to the same time point as wildtype nucleosome arrays ($* = p < 0.05$, $** = p < 0.01$, $*** = p < 0.001$). SDS-PAGE gels show representative ubiquitylation assays of indicated nucleosome arrays. The upper panel displays in-gel fluorescence of fluorogenic H2A, and the lower panel shows the same gels stained with Coomassie blue (CM).

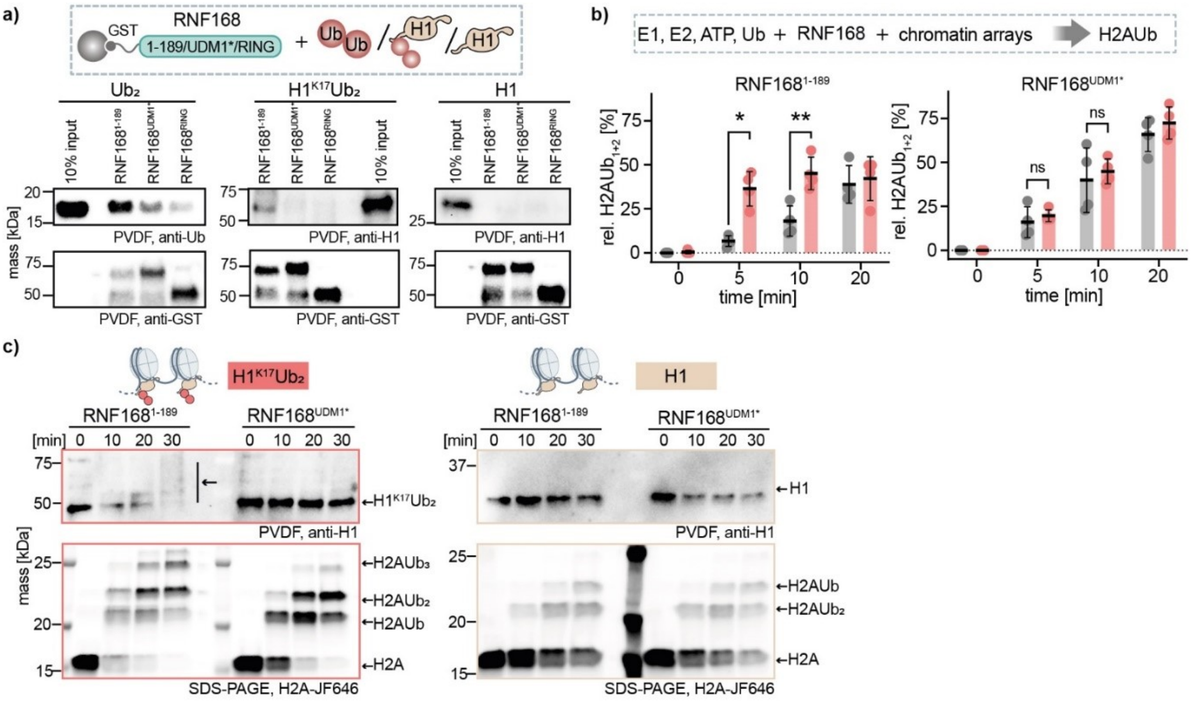

**Figure S7.** a) Western blot analysis of RNF168 variant pull-downs of $Ub_2^{N3}$ (α-Ub), $H1^{K17}Ub_2$ (α-H1) and H1 (α-H1). RNF168 loading controls (α-GST) are shown in the lower panel. b) Quantification of H2A in-gel fluorescence intensities corresponding to the sum of the H2AUb and $H2AUb_2$ bands normalized to H2A levels at t = 0 min. Data represent the mean and standard deviation of four independent replicates with individual data points indicated. Significance is indicated for the time points t = 5 min and t = 10 min ($*$ = $p < 0.05$, $**$ = $p < 0.01$, $***$ = $p < 0.001$). c) Western blot analysis (α-H1, top) of ubiquitylation assays by $RNF168^{1\text{-}189}$ and $RNF168^{UDM1}$ in presence of $H1^{K17}Ub_2$ (left) or H1 (right). In-gel fluorescence (below) shows the corresponding ubiquitylation of H2A. (PVDF: polyvinylidene fluoride membrane)

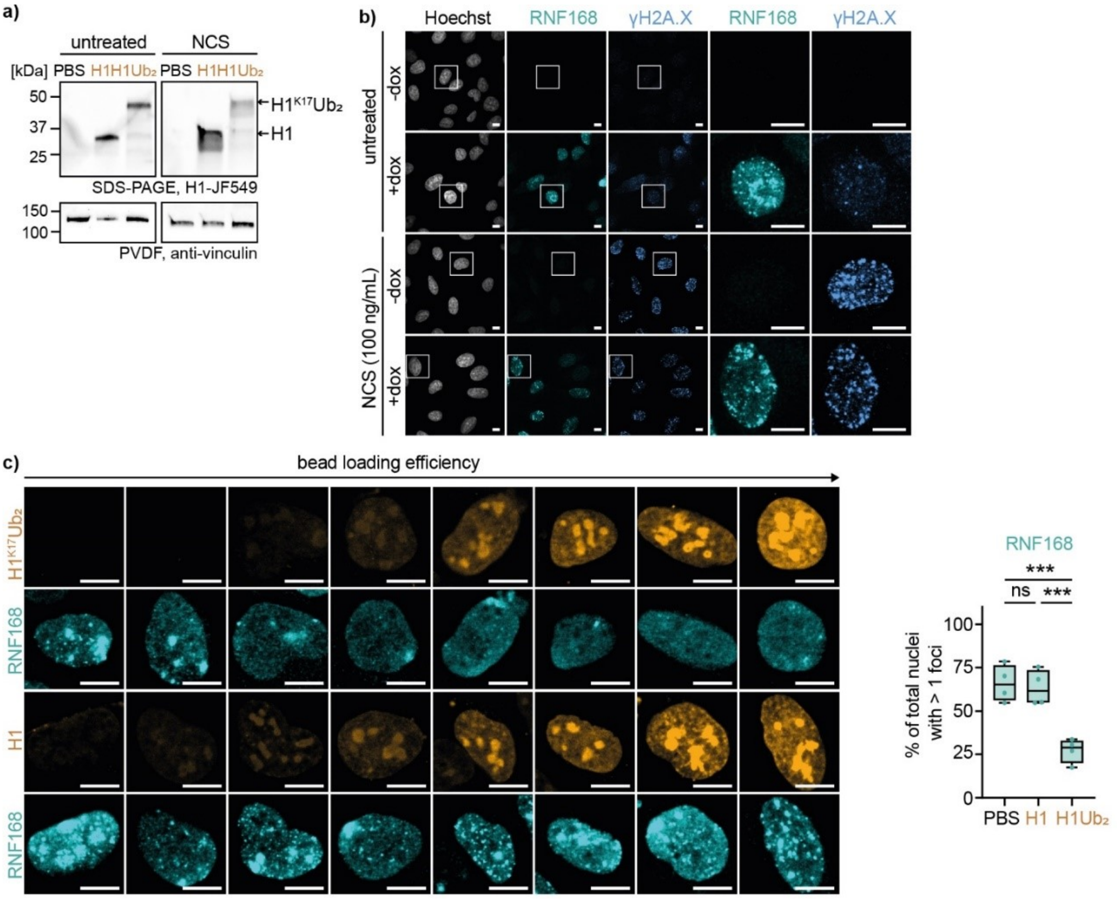


**Figure S8.** a) FLAG-RNF168 expression was induced in Flp-In U-2 OS cells by doxycycline (dox, 1 µg/mL, 24 h). Then, cells treated with vehicle or NCS (100 ng/mL for 1 h) before and after bead-loading of PBS or fluorescently labeled H1 or $H1^{K17}Ub_2$ and analyzed by SDS-PAGE and Western Blot (α-vinculin, AB155120, Abcam) (PVDF: polyvinylidene fluoride). b) Dox-induced U-2 OS Flp-In FLAG-RNF168 cells were treated either with vehicle (0.02% 20 mM 2-(*N*-morpholino)ethanesulfonic acid (MES)) or neocarzinostatin (NCS) (100 ng/mL for 2 h). FLAG-RNF168 (cyan) and γH2AX (blue) were visualized via immunofluorescence. c) (related to Figure 4c) Dox-induced U-2 OS Flp-In FLAG-RNF168 cells were bead-loaded with PBS or fluorescently labeled H1 or $H1^{K17}Ub_2$. FLAG-RNF168 (cyan) was visualized via immunofluorescence. Representative nuclei displayed varying bead-loading efficiencies within the same experiment. While RNF168 foci formation did not depend on H1, it concentration-dependently declined with increasing levels of $H1^{K17}Ub_2$ (scale bar 10 µm). Proportion of nuclei containing more than one RNF168 focus quantified on the right.

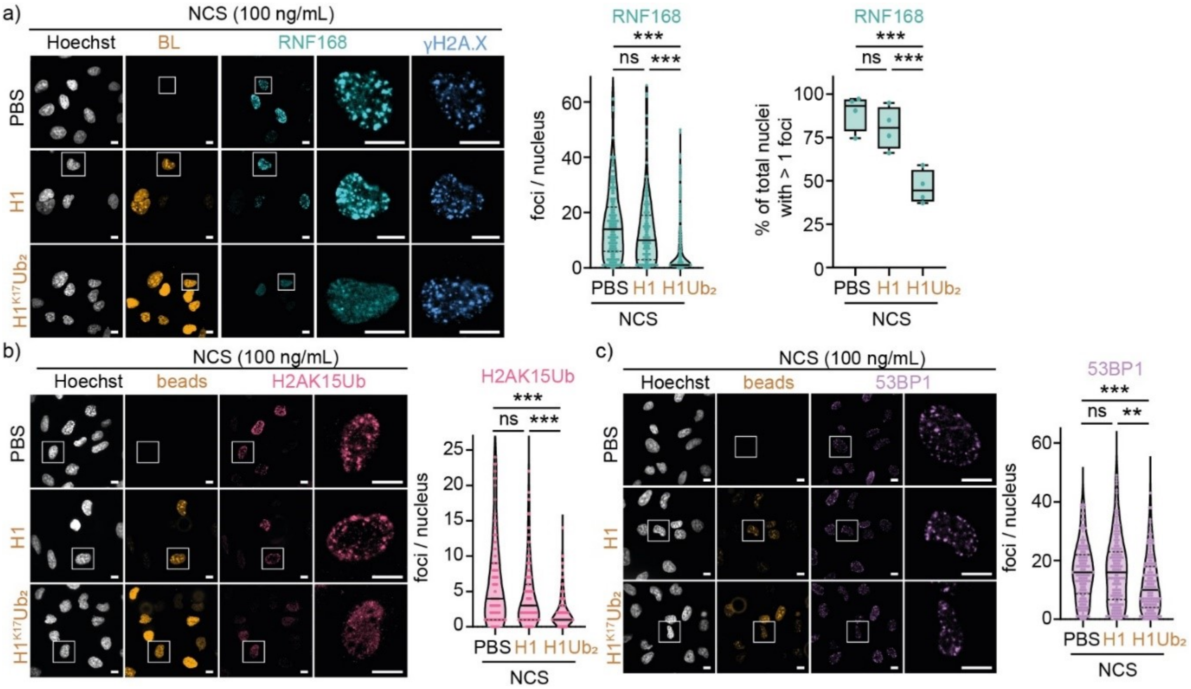


**Figure S9.** a) Dox-induced U-2 OS Flp-In FLAG-RNF168 cells were treated with NCS (100 ng/mL) for 1 h before and after bead-loading of fluorescently labeled H1 or H1$^{K17}$Ub$_2$. FLAG-RNF168 (cyan) and $\gamma$H2A.X (blue) were visualized via immunofluorescence (left). Quantification of RNF168 foci and proportion of nuclei containing more than one RNF168 focus (right). Data represent four replicates with $n > 180$ nuclei per condition analyzed. b) Dox-induced U-2 OS Flp-In FLAG-RNF168 cells were bead-loaded with PBS or fluorescently labeled H1 or H1$^{K17}$Ub$_2$. H2AK15Ub (red) was visualized via immunofluorescence. Quantification of H2AK15Ub foci is shown on the right. Data represent two independent replicates with $n > 110$ nuclei per condition analyzed. c) Dox-induced U-2 OS Flp-In FLAG-RNF168 cells were bead-loaded with PBS or fluorescently labeled H1 or H1$^{K17}$Ub$_2$. 53BP1 (purple) was visualized via immunofluorescence. Quantification of 53BP1 foci (is shown on the right). Data represent two independent replicates with $n > 110$ nuclei per condition analyzed. ($* = p < 0.05$, $** = p < 0.01$, $*** = p < 0.001$, scale bar 10 µm).

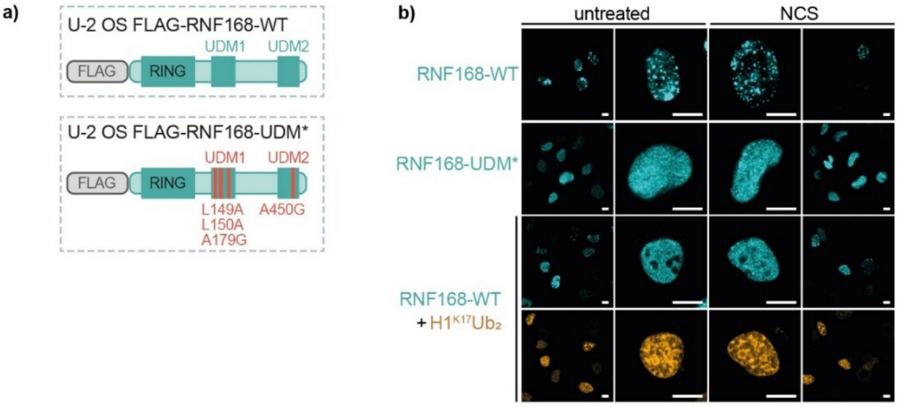


**Figure S10.** a) U-2 OS Flp-In cells expressing either FLAG-RNF168-WT or the ubiquitin-binding deficient FLAG-RNF168-UDM* carrying four point mutations in UDM1 and UDM2[7]. b) RNF168-UBD* failed to form foci even upon DNA damaging treatment by NCS, mimicked by RNF168-WT cells bead-loaded with H1$^{K17}$Ub$_2$ (scale bar 10 µm).
